# Enhanced Neutralization of Japanese Encephalitis Virus Using an Engineered VL Double-Domain

**DOI:** 10.1101/2025.09.03.673967

**Authors:** Mario Marweslie, Ika Agus Rini, Misae kiba, Ramadhani Qurrota Ayun, Taek-Kyun Lee, Sukchan Lee

## Abstract

Japanese encephalitis virus (JEV) is an emerging virus responsible for thousands of deaths in Asia; however, an effective treatment has not yet been discovered. JEV envelope (E) proteins play important roles in viral survival and infection. In this study, we isolated scFv- and VL-targeting E proteins using phage display biopanning. The isolated VL was engineered to form VL double domains to increase its neutralization activity. Several candidates showed binding abilities, whereas only the DE2 VL double-domain treatment showed neutralizing activity against JEV. DE2 was also highly specific to JEV and did not bind to other flaviviruses. Immunocytochemistry results showed that DE2 was co-localized with the endosome 4 h post-infection. Thus, the mechanism of DE2 neutralization involves blocking the E protein from fusing with the endosomal membrane, resulting in the unrelease of the viral genome into the cytoplasm. Docking analysis showed that DE2 was bound to three domains, especially DIII, of the JEV E protein. In contrast, other scFv that were unable to bind to DIII showed no or low neutralizing activity. DE2 was also predicted to neutralize all the JEV genotypes. In conclusion, a combined in vitro and in silico study showed the high potential of DE2 to neutralize JEV infections, which may be applied in future therapeutic systems.

**Importance:** Japanese encephalitis virus (JEV) is a serious mosquito-borne disease that causes brain infections and deaths across Asia. However, there is still no specific treatment once infection occurs. In this study, we developed a new type of antibody fragment, called a VL double-domain, that can block the virus. Among several candidates, DE2 showed a strong ability to stop JEV from infecting cells, while others could bind to the virus but did not block infection. Our findings reveal that DE2 works by attaching to a key region of the virus, preventing it from releasing its genetic material. Moreover, DE2 is predicted to work against all known JEV strains, highlighting its potential as a universal treatment option. This research provides an important step toward developing new antiviral therapies for JEV and possibly for other viruses with similar infection mechanisms.

## 1. Introduction

The Japanese encephalitis virus (JEV) is the leading cause of viral encephalitis worldwide [1,2]. An estimated 100.000 cases of JEV infection occur each year in Asia, with two out of three classified as severe [3,4]. JEV is a mosquito-borne virus that belongs to the flavivirus family. It spreads zoonotically, with mosquitoes as vectors, birds and pigs as amplifying hosts, and other vertebrates such as horses and humans as dead-end hosts [5,6]. Currently, there is no effective treatment for JEV infections other than vaccines for prevention [7,8]. Therefore, developing an effective treatment is essential to control diseases caused by JEV infections [9].

JEV is an enveloped virus with a virion size of 50 nm that consists of positive-sense single-stranded RNA with a total genome size of approximately 11 kb [10]. The envelope protein (E) of JEV is located on the surface of the virus and plays an important role in host cell interactions, including receptor binding, transduction, and membrane fusion. The E protein of JEV has three domains: domain I (central β-barrel), domain II (fusion loop), and domain III (immunoglobulin-like structure). In its mature form, the E protein forms a flattened dimeric structure [10–12]. Many monoclonal antibodies that show strong neutralizing activity against flaviviruses target the E protein. Therefore, the E protein is a crucial viral antigen that induces and serves as a target of neutralizing antibodies [10,13,14].

Recombinant antibodies are widely used in diagnostic and therapeutic research [15,16]. One commonly used recombinant antibody is the single-chain variable fragment (scFv), which consists of variable regions of the heavy chain (VH) and light chain (VL). ScFv was commonly applied to detect and neutralize various viruses. Because of its small molecular size, scFv can be easily expressed and modified [17,18]. Phage display is commonly applied for antibody screening. This method significantly reduces the time and cost of antibody screening [18,19]. In previous studies, some scFvs have been engineered or modified to improve their antigen-binding affinity, stability, and solubility [20–22]. Furthermore, single-domain VL and scFv fragments were modified to be bivalent, which enhanced their viral binding [20,23].

In this study, we screened neutralizing antibodies with phage display biopanning against JEV. scFv was also engineered to enhance its neutralizing activity against JEV by changing the domain into VL-VL (VL double domain) form. To further elucidate the neutralizing mechanism of action, a docking analysis was performed to predict the binding site of scFv to JEV, which provided the structural basis for its neutralizing activity.

## 2. Materials and Methods

### 2.1 Cells and viruses

Vero E6 cells were maintained in Dulbecco’s modified Eagle’s medium (DMEM) (GenDEPOT) supplemented with 10% heat-inactivated fetal bovine serum (FBS) (GenDEPOT) and 1% Antibiotic-Antimycotic solution (HyClone). Cells were cultured in a 37°C incubator with 5% CO_2_.

JEV genotype III (KBPV-VR-27) and Dengue virus serotype 2 (KBPV-VR-29) were obtained from the Korea Bank for Pathogenic Viruses (KBPV), Zika virus (NCCP N0. 43280) obtained from the Korea Disease Control and Prevention Agency. All Viruses propagated in Vero E6 cells using Eagle’s minimal essential medium (MEM) (Gibco). Virus was harvested at 6–7 days post-incubation. Supernatants were clarified by centrifugation (1,000 × g, 5 min), followed by ultracentrifugation (35,000 rpm, 2.5 h). Pelleted virus was resuspended in PBS, filtered (0.22 µm), and stored at –80°C. [24].

### 2.2 ScFv selection through biopanning and ELISA

Biopanning was performed as previously described. Briefly, 96-well plates were coated with JEV (10⁴ PFU/mL) or PBS (control) overnight at 4°C, washed with TBST (0.1% Tween 20 in Tris-buffered saline), and blocked with 5% BSA. The Tomlinson I+J scFv phage library was pre-incubated in control wells before transfer to JEV-coated wells. Bound phages were eluted with 100 mM triethylamine, neutralized with 1 M Tris-HCl (pH 7.0), and amplified in XL-1 Blue on LB agar with tetracycline and ampicillin. Five rounds of biopanning were performed. [20,23,25,26].

To identify specific candidates, ELISA was performed in 96-well plates coated with JEV (overnight, 4°C), blocked, and incubated with phage clone supernatants or buffer (control). After washing, HRP-conjugated anti-M13 antibody (1:1000; Sino Biological) was added, followed by TMB substrate. The reaction was stopped with 1 N H₂SO₄, and absorbance was measured at 450 nm. Selected scFv phages were cloned and analyzed using IgBlast (NCBI) [20,23,25].

### 2.3 ScFv expression and purification

Selected scFv genes were cloned into the pIG20 vector and expressed in *Escherichia coli* BL21 (DE3) pLysE. Protein expression was induced with 1 mM IPTG at 20°C or 25°C overnight, and large-scale production was performed under optimal conditions. scFvs were purified from the supernatant using Capto L resin (Cytiva). Purity was assessed by SDS-PAGE followed by Coomassie blue staining and destaining [20,23,25].

### 2.4 Binding ability test of scFv candidates by ELISA

The JEV in coating buffer (carbonate/bicarbonate buffer, pH 9.6) was coated on 96-well plates overnight at 4°C. Blocking buffer (5% skim milk in TBST) was added to the plate and incubated for 2 h at 25°C. After five washes, scFv (100 µg/mL) was added and incubated for 2 h at 37°C. This was followed by incubation with Chicken anti-Protein A (1:10.000) (Sigma) for 1 h at 37°C. After the final washing, TMB solution was added, and the absorbance value at 450 nm was measured using a spectrophotometer [20,23,25].

### 2.5 Neutralizing assay

The JEV (MOI 0.01) was neutralized by 200µg/μL antibodies for 1 h at 37°C. The virus-antibody mixture was then incubated with Vero E6 cells for 1 h at 37°C. The cells were then cultured in MEM supplemented with 10% FBS and 1% antibiotic-antimycotic solution. The cells were harvested after 48 h post-infection (hpi) and stored at −20°C for further RNA extraction. Neutralization assays for Zika and Dengue viruses were performed using the same methods [27,28].

### 2.6 RT-qPCR analysis of gene expression level

Total RNA was extracted using TRI reagent (MRC), and 1 µg was reverse-transcribed with the PrimeScript 1st Strand cDNA Synthesis Kit (Takara). qRT-PCR was performed using TB Green® Premix Ex Taq™ II (Takara) on a Rotor-Gene Q system (Qiagen) with primers listed in Table S1. GAPDH served as a control for relative expression analysis [27,28]. Viral attachment and internalization after DE2 treatment were analyzed by RT-qPCR. For attachment, cells were incubated with the JEV-DE2 mixture at 4°C for 1 h, washed with cold PBS, and RNA was extracted. For internalization, cells were incubated with JEV-DE2 at 4°C for 1 h, washed, supplied with fresh medium, and incubated at 37°C for 1 h before RNA extraction and analysis [29,30]

### 2.7 Western Blotting

Total protein was extracted using PRO-PREP™ (Intron Biotechnology). Samples (30 µL) were separated by SDS-PAGE, transferred to nitrocellulose membranes, and probed with anti-JEV NS3 (1:5000; GeneTex) and anti-GAPDH (1:2000; Cell Signaling) primary antibodies, followed by HRP-conjugated anti-rabbit (1:10,000) and anti-mouse (1:5000; Cell Signaling) secondary antibodies. Bands were visualized with the iBright™ CL1500 Imaging System (Thermo Fisher), and relative intensity was quantified using ImageJ [27,28].

### 2.8 Plaque reduction assay

The supernatant harvested from the neutralizing procedure was used to infect Vero E6 cells. After 1 h incubation at 37°C, the cells were washed and overlaid with 1% Seaplaque agarose containing MEM with 1% PSA antibiotic. The cells were fixed with 5% formaldehyde and stained with 0.5% crystal violet after 7 days of incubation at 37°C [24,27].

### 2.9 Immunocytochemistry (ICC)

Vero E6 cells were cultured in 8-well chambers, infected with JEV, and treated with antibodies as described. Cells were fixed with methanol, permeabilized, and blocked before incubation with primary antibodies against JEV NS3 (1:750; GeneTex), Protein A (1:750; Exalpha), and EEA1 (1:750; Abcam) overnight at 4°C. Alexa Fluor– conjugated secondary antibodies (1:1000; Abcam) were applied for 1 h at 25°C, and nuclei were counterstained with DAPI (LSbio). Images were acquired using an LSM 900 confocal microscope (Zeiss). Relative intensity was calculated as viral (green) to nuclear (blue) signal ratio and normalized to untreated controls. [27,28].

### 2.10 In silico analysis

Molecular docking was used to predict the binding interactions between scFv and JEV envelope protein. The scFv structure was modeled using the Swiss Model server and validated with the PROCHECK server using a Ramachandran Plot. The scFv model with less than 90% of the residues in the favored region was refined using the GalaxyWEB server. The JEV E protein crystal structure was retrieved from the Protein Data Bank (PDB 3P54) [11]. scFv and JEV E proteins were docked using HADDOCK 2.4. The best model was selected based on HADDOCK and Z-values. The docking model was then analyzed using LigPlot^+^ to generate the scFv-JEV E protein interaction, as shown by hydrogen bonds and hydrophobic contacts. The docking complex was also visualized using UCSF Chimera. Alignment of envelope protein sequences was visualized using ESPript 3.0 [31].

## 3. Results

### 3.1 Screening, modification, and characterization of JEV-specific scFv

To identify JEV-specific antibodies, five rounds of biopanning were performed using whole viral particles as antigens (Figure 1A). The antigen-binding absorbance was high in the fifth round, whereas the first four rounds showed low absorbance. This was due to the presence of many scFv with low or no binding to the antigen, which resulted in lower average absorbance during the earlier rounds. Scfv phages that demonstrated binding ability to the antigen were collected in each round. The VH and VL sequences were cloned into the pIG20 vector for further cloning and expression (Figure 1B).

**Figure 1.**
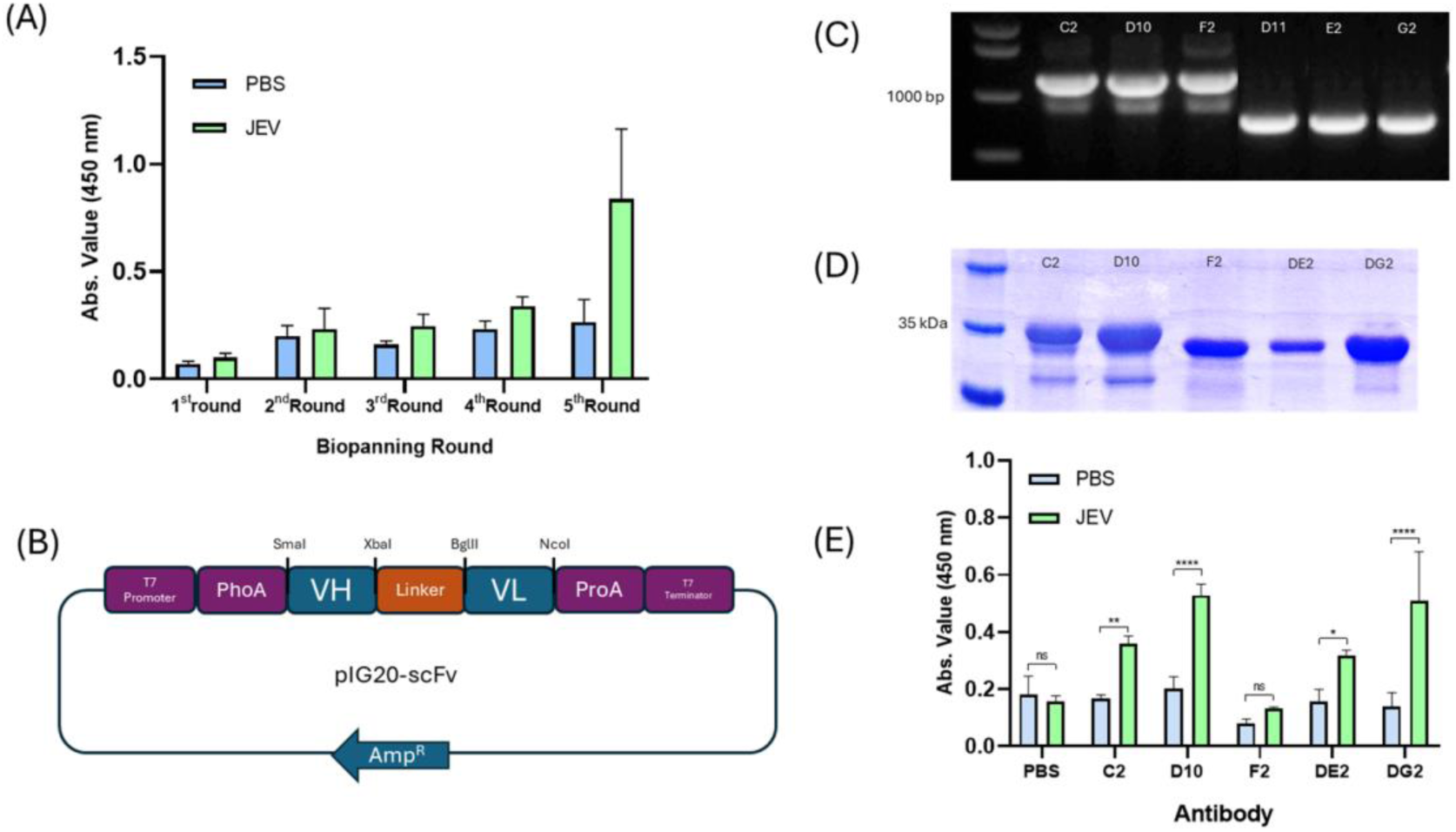
Isolation and characterization of scFvs using biopanning. (A) Biopanning average absorbance from each round. Five rounds of biopanning were conducted, and scFvs were collected each round. (B) Schematic diagram of vector for scFv cloning and expression. scFv sequences obtained from biopanning were cut and ligated to pIG20. (C) Verification of insertion and size of scFv in pIG20 using PCR. Three scFv sizes are approximately 900 bp, and 3 VL single domains are approximately 400 bp. (D) Size verification of purified scFv. All scFv and VL double domain sizes are approximately 35 kDa. (E) ELISA analysis of scFv binding to JEV. All antibodies can bind to JEV, except F2 scFv.

Three scFv (C2, D10, F2) and three single-domain VL (D11, E2, G2) clones were identified as anti-JEV candidates. Complementarity-determining regions (CDR) sequence analysis showed identical CDR1 regions among scFv clones, with variations in CDR2 and CDR3 (Table 1). The antibodies were cloned as approximately 900 bp scFv and 400 bp VL domains (Figure 1C). VL domains were further engineered into double domains (DE2, DG2) (Figure S1A). The CDR regions were analyzed for CDR composition (Table S2) and successfully expressed (Figure S1B).

**Table 1.**
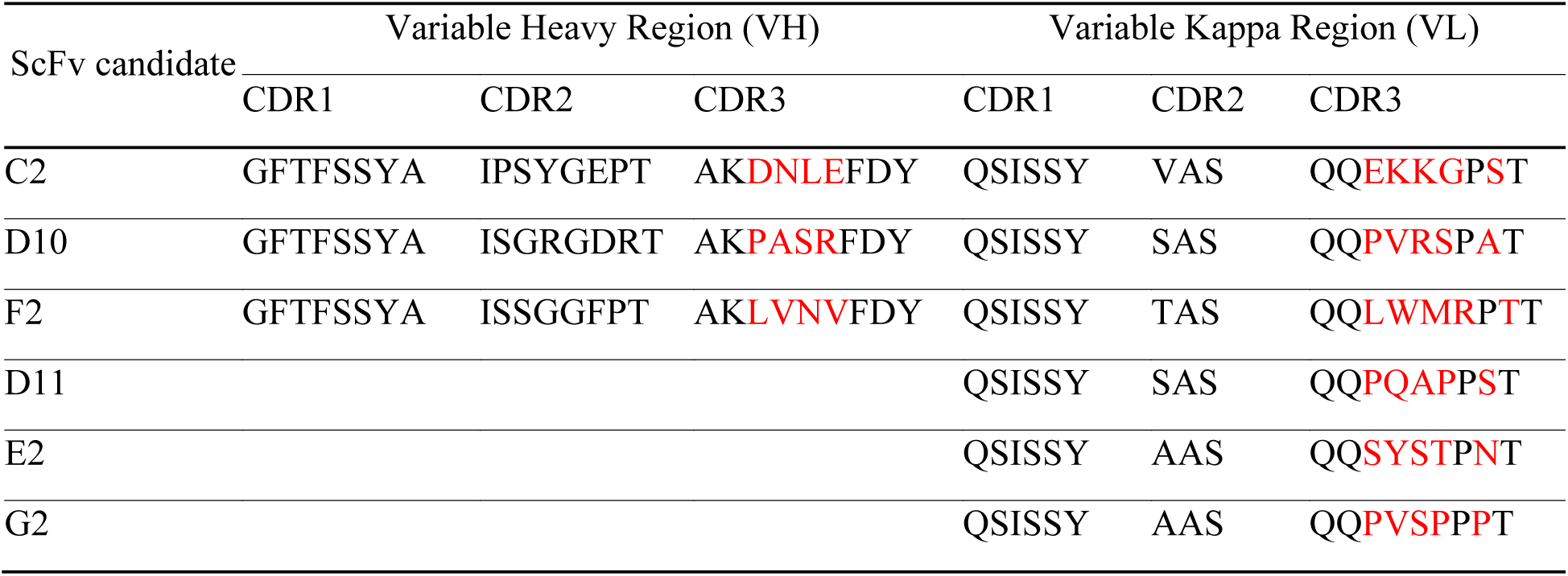
CDR analysis of scFv candidates obtained from biopanning.

Small-scale expression was conducted at two different temperatures after induction to determine the optimal expression conditions (Figure S2). Subsequently, the scFv was expressed on a large scale and purified using a strong-affinity resin against the kappa chain. Purified scFv was confirmed using SDS-PAGE (Figure 1D). Each scFv showed a different yield; however, the highest yield was obtained with DE2 (6.5 mg/L) (Table S3). An ELISA was performed to evaluate the binding ability of scFv clones to JEV (Figure 1E). C2, D10, DE2, and DG2 demonstrated binding abilities to JEV, as indicated by higher absorbance values, whereas F2 showed low absorbance, indicating no binding to JEV.

### 3.2 Neutralizing activity of scFv against JEV

Four candidate antibodies with binding abilities were further utilized to determine their neutralization activity against JEV. RT-qPCR was used to measure the relative expression of the JEV E and NS1 genes (Figure 2A). Based on the relative expression of the E and NS1 genes, DE2 and DG2 exhibited neutralizing activities of approximately 50% and 20%, respectively. In contrast, C2 and D10 did not show any neutralizing activity compared to the positive control.

**Figure 2.**
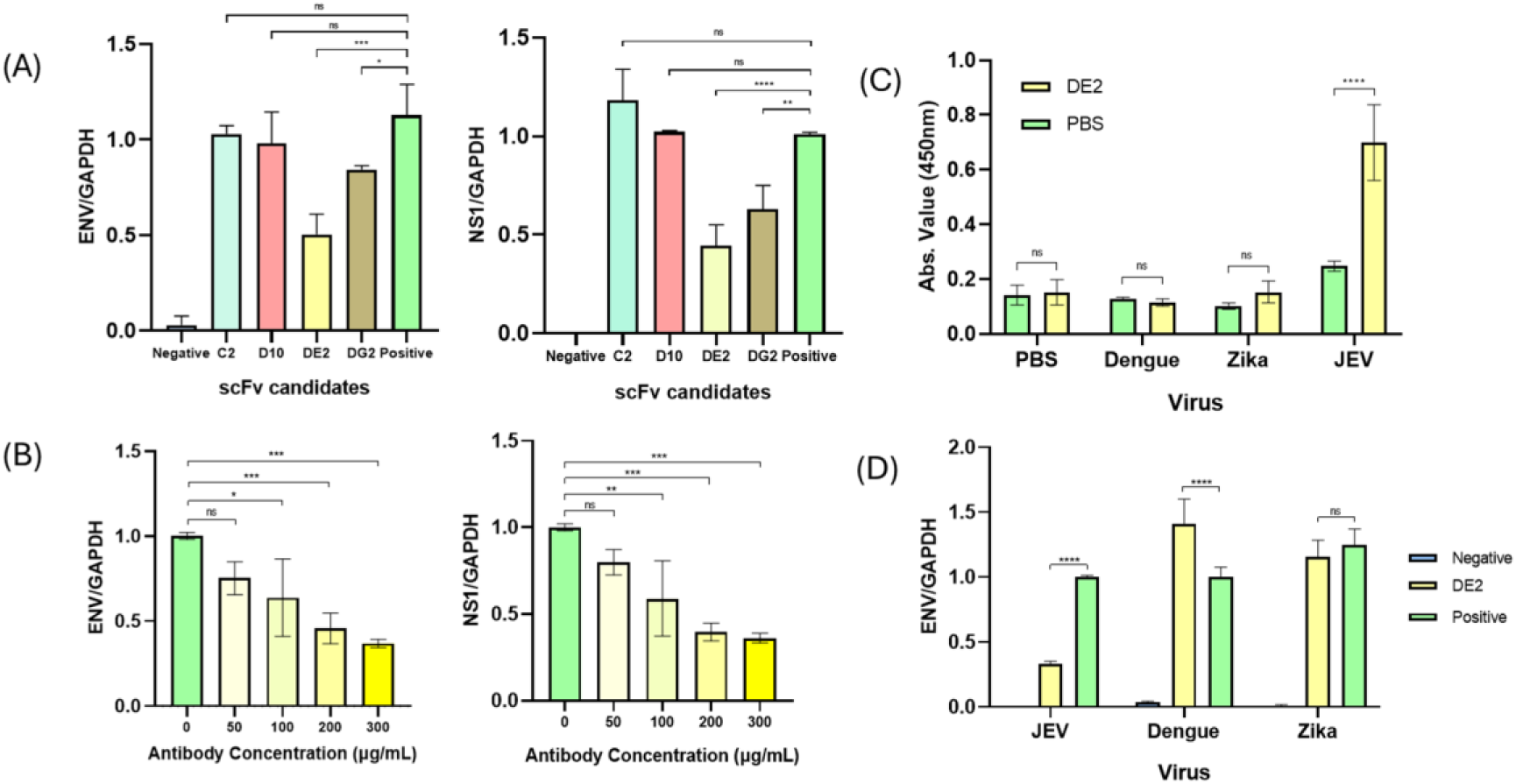
Characterization of scFv neutralizing ability against JEV. (A) Expression of JEV measured by the ENV and NS1 genes from each antibody candidate. DE2 and DG2 have lower JEV levels than the positive control. (B) Measurement of JEV relative expression with various concentrations of DE2. The optimum concentration of DE2 was 200 µg/mL. (C) Specificity test of DE2 against dengue and Zika using ELISA (D) Neutralization assay of DE2 against dengue and Zika virus. DE2 binds specifically and only neutralizes JEV. All virus-specific gene expression was normalized to GAPDH expression.

The most potent neutralizing antibody, DE2, was evaluated for cytotoxicity at various concentrations (Figure S3). The result showed that there is no reduction in cell viability with DE2 up to 300 µg/mL. Various concentrations of DE2 were used to neutralize JEV (Figure 2B). The optimum concentration to neutralize JEV with 50% inhibition was 200 µg/mL.

DE2 was also compared to its single domains (D10 VL and E2 VL) to determine whether the single domain alone had the same neutralization activity (Figure S4). However, only DE2 possessed a neutralization ability, whereas the single domain did not inhibit JEV infection. To check the specificity of DE2, ELISA and neutralization assays were conducted using other flaviviruses, such as dengue and Zika virus (Figure 2C, 2D). The results showed that DE2 bound only to JEV. The neutralization assay also showed that DE2 inhibited viral propagation in JEV, but not in dengue and Zika, showing that DE2 was highly specific to JEV.

Viral titer in the supernatant after neutralization was measured in the plaque reduction assay. DE2 decreased the viral titer by 10-fold compared to the positive control (Figure 3A). The expression level of JEV NS3 protein was also decreased in DE2 treatment by approximately 50% compared to the positive control, as determined by western blotting analysis (Figure 3B). Immunocytochemistry results showed that the JEV signal was reduced by approximately 63% after DE2 treatment compared to that in the positive control (Figure 3C).

**Figure 3.**
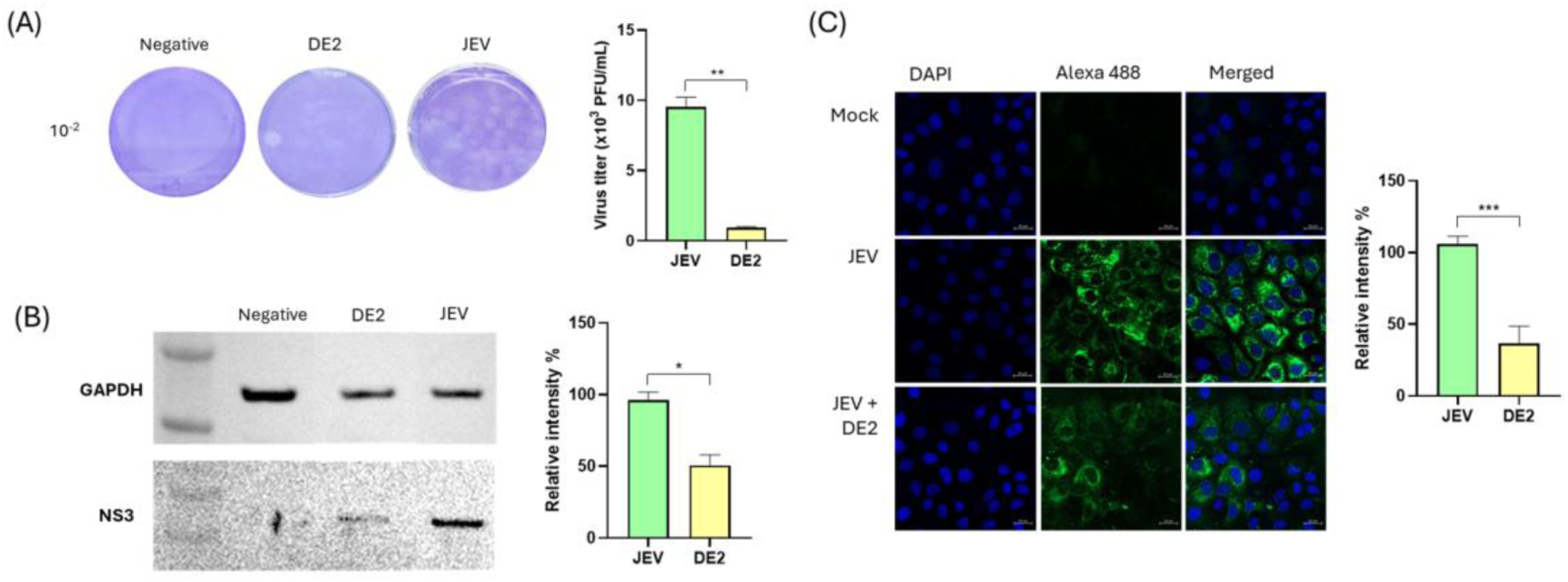
Inhibition of JEV by DE2 *in vitro*. (A) The JEV infectious titer was measured using a plaque reduction assay. (B) The NS3 protein of JEV was examined using western blotting. The bands’ intensity was determined using ImageJ. (C) JEV visualization after neutralization at 48 hpi by LCM. JEV signals (in green, Alexa 488) were calculated using ZEISS LSM software and normalized to the cell’s nucleus signal (in blue, DAPI) to evaluate the JEV titer treated with DE2.

We examined the immune response of Vero E6 cells during JEV infection following DE2 treatment (Figure S5). Cytokines (IL-6, IL-18, and TNF-) and chemokines (CCL5 and CXCL10) were significantly downregulated in the DE2-treated group compared to the positive control. Receptors involved in the inflammatory response (TLR-7 and TLR-3) also showed lower expression levels in response to DE2 treatment. These results show that DE2 treatment protected cells from JEV infection.

### 3.3 Neutralizing activity of scFv against JEV

To assess the mechanism of action of DE2 in neutralizing the virus, attachment assays were conducted to determine whether the virus was blocked from binding to the host cell receptors (Figure 4A,B). The results showed that DE2 blocked viral attachment to the host cells by approximately 20%. Even at higher concentrations, it did not increase viral attachment blocking, suggesting that the DE2 mechanism of action does not block attachment to the host cell. Internalization assays were also conducted to determine whether the virus was blocked during internalization into the host cell membrane; however, DE2 did not block the virus from entering the cell membrane (Figure S6A).

**Figure 4.**
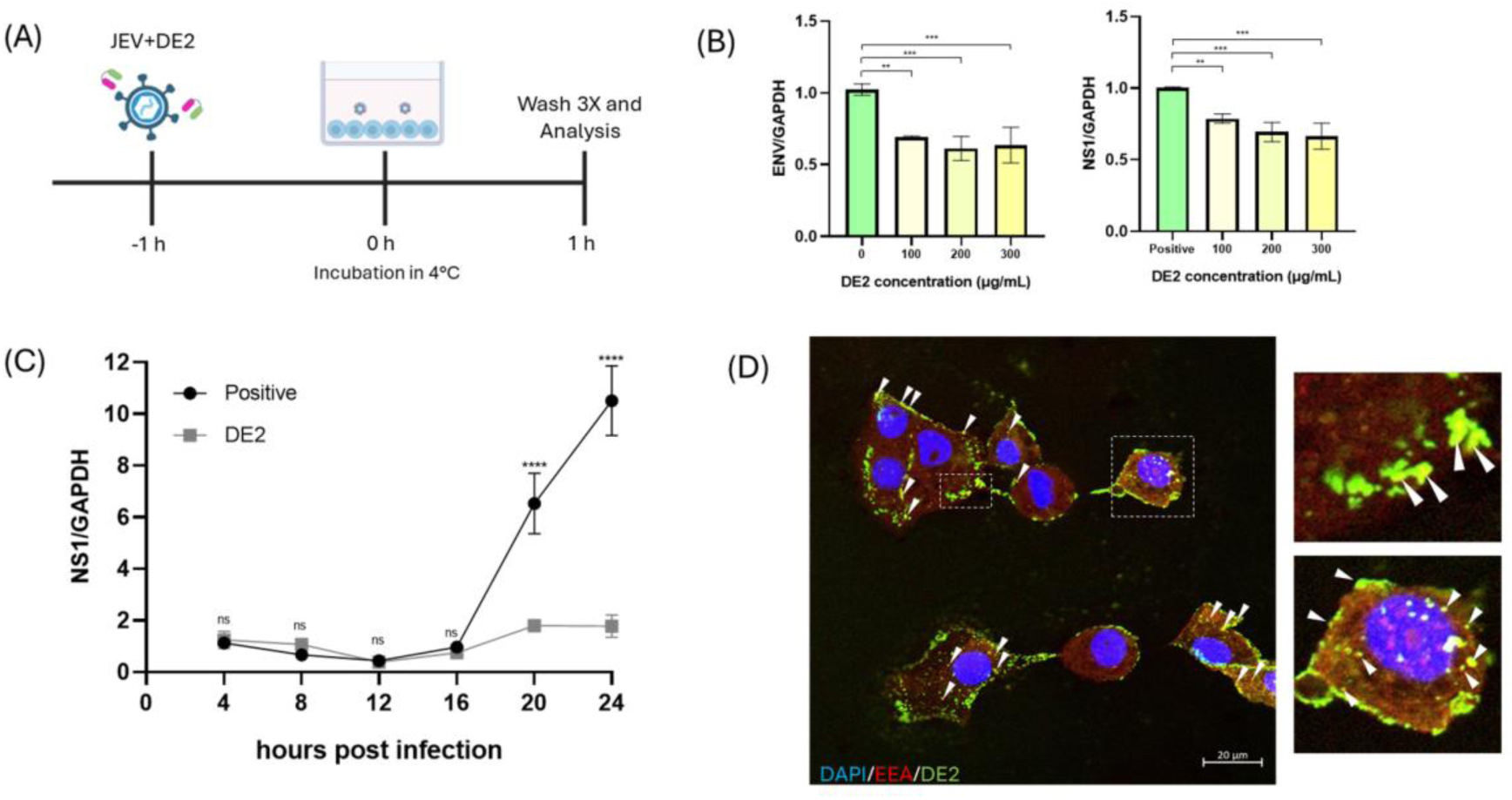
DE2 neutralizes JEV by inhibiting attachment and endosome fusion. (A) Schematic diagram of the attachment assay. Cells were incubated with the antibody-virus mixture at 4°C to measure viral titer on the cell surface. (A) Relative expression of ENV and NS1 genes of JEV on the cell surface detected by real-time PCR. (B) Relative expression of NS1 genes JEV in a time-dependent manner. JEV was blocked after 20 hpi. (C) JEV-bound DE2 was internalized into the cell and localized to the endosome. DE2 (in green, Alexa 488) and EEA-1 (in red, Alexa 647) signals were visualized after 4 hpi (MOI 0.1) using LSM. Images inside the squares with the dashes are shown at higher magnification.

We quantified the viral genes until 24-hpi (Figure 4C). During 0-16 pi, the viral amount between the untreated and treated with DE2 remains the same. The neutralization effect began to appear after 20 hpi. At 24 hpi, the DE2 treatment exhibited a viral titer similar to that at 4 hpi, whereas in the positive control, the viral titer increased 5.9-fold from 4 hpi.

Time-dependent assay analysis showed that the neutralizing process did not occur during viral attachment or entry, suggesting that DE2 blocks the virus when it fuses with the endosome. ICC was conducted to determine viral and DE2 localization in host cells (Figures 4D and S6B). This indicated that DE2 was bound to JEV and was localized in the endosome. These findings suggest that the neutralizing mechanism of action of DE2 blocks the release of the genome from the endosome.

To elucidate the capability of DE2 to neutralize JEV, a docking simulation of DE2 and the JEV E protein was conducted. The JEV E protein was retrieved from the PDB, as verified by its crystal structure. The predicted antibody model was obtained from the Swiss Model and validated using the PROCHECK server. Some antibody models were optimized to yield high-quality models with more than 90% of the residues in the core region, as evidenced by Ramachandran plots. The Ramachandran plot of the DE2 model showed an accuracy of 86.4% in the favored region. After optimization, 90.6% of the residues were in the favored region, 8.4% in the allowed region, and 1% in the disallowed region (Figure S7A).

Subsequently, molecular docking was employed to analyze the interaction between the JEV E protein and the antibody. The best docking model exhibited the highest HADDOCK and lowest Z-scores (Table S4). The best docking model was visualized using UCSF Chimera and processed with LigPlot+ to identify the amino acids involved in the interactions (Figure 5 and S7B).

**Figure 5.**
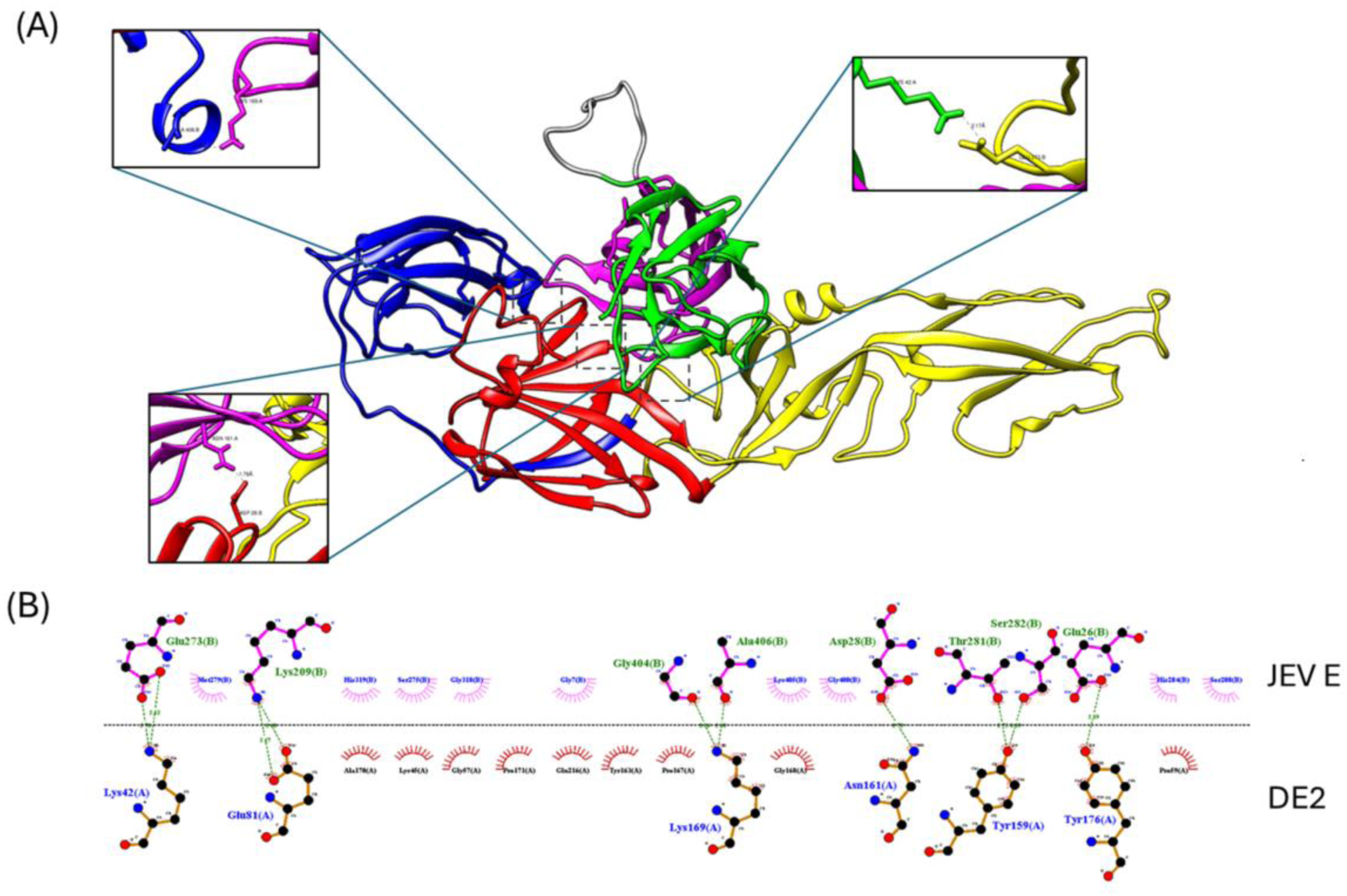
Molecular interaction between DE2 with JEV envelope protein. (A) Docking analysis to visualize the 3D structure. In the enlarged image, the dashes between proteins showed the hydrogen length. DI, DII, and DIII regions are colored in red, yellow, and blue, respectively. The first VL is colored green, and the second VL is colored pink. (B) Specific interaction analysis for visualization of the 2D structure. Green dashed lines indicate hydrogen bonds, and red spoked arcs indicate hydrophobic interactions.

Based on the docking models, DE2 bound to all three domains of the E protein, similar to DG2 (Table S5). D10 scFv bound only to domain II, whereas C2 scFv bound only to domains I and II. We further examined whether the binding sites of DE2 in the E protein were conserved. The JEV E proteins from genotypes I–V were retrieved and aligned using the protein sequence model. The results showed that the epitopes were highly conserved in all genotypes, with epitope sequence similarities between GI, GIII, GIV (94%), and GII; GIV being 88%. Compared to other flaviviral E proteins, Zika and DENV2 had only 58% and 41% epitope sequence similarity, respectively.

## 4. Discussions

JEV is considered to be an emerging and re-emerging virus owing to its various genotypes that cause infections in 24 countries across Asia and Oceania. Although JEV infection is vaccine-preventable, the current specific drugs for JEV are still limited [32–34]. scFvs are a well-established antibody format that have been widely used for viral detection and neutralization of diverse viruses [35–37]. In this study, we successfully generated neutralizing antibodies against JEV using phage display with the human scFv library (Tomlinson I + J), resulting in both scFv and VL formats [26].

We engineered a single VL domain to form a VL double domain (DE2 VL). The VL single domain is relatively unstable and has a lower binding ability than the double-domain form [38–40]. In many studies, scFv was engineered to increase its functionality, whether by tagging with a peptide or chemical, or by changing the scFv construct, such as making it bivalent [20,41–43]. In a previous study, the single VL domain was converted to a bivalent form, which showed increased neutralization activity [23]. In this study, instead of combining the same VL, we combined two different VL to target different antigen binding sites. Engineered DE2 showed the highest yield of expressed proteins, indicating its better stability and solubility. Compared with the single domains (VL D10 and VL E2) and the other scFv, DE2 showed the best neutralizing activity against JEV.

We found that DE2 binds to JEV but cannot completely block the virus attachment to the cell. Since JEV can replicate in many cell types, it likely employs various receptors for entering the cell [14,44]. Although DE2 is used at high concentrations to potentially bind and cover the virus surface, some virions still can enter the cell, suggesting that blocking attachment is not the mechanism of action.

After entry into the cell, the virus remains in the endosome before fusing with the endosome and releasing its genome into the cytoplasm. During fusion, the JEV envelope protein disassociates and forms a trimeric structure [6,10,11]. We observed that, in the presence of DE2, the viral titer remained unchanged until 24 hpi, whereas without DE2 showed clear viral propagation. This indicates that the virus was able to enter the cell but was unable to propagate, further supporting that DE2 does not block the viral attachment.

Furthermore, DE2 localized to the endosome after 4-hpi. scFv is normally unable to enter the cell unless engineered or bound to substances that can enter the cell [45]. Although DE2 cannot enter the cell under normal conditions, it was able to enter the cell by binding to the virus. Because DE2 was localized in the endosome, we proposed that its neutralization mechanism involves inhibition of endosomal fusion, thereby preventing the release of the JEV genome into the cytoplasm (Figure 6).

**Figure 6.**
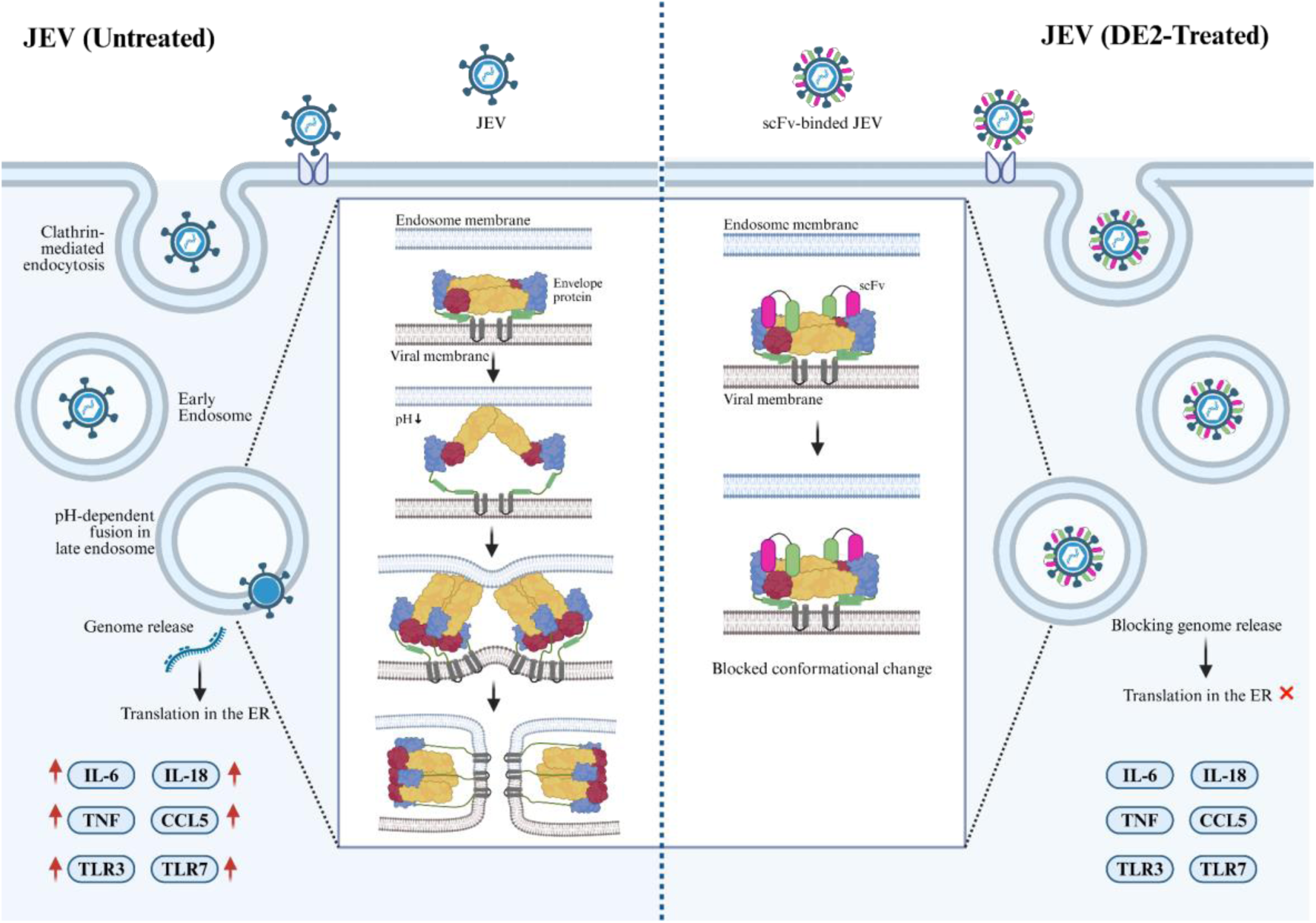
Proposed model of the neutralizing mechanism of action of DE2. DE2-bound JEV enters the cell in the endosome by endocytosis. The E protein undergoes conformational change owing to low pH and fusion with the endosome membrane, resulting in genome release and upregulation of the immune response. However, the DE2-bound E protein is unable to change its conformation. As a result, the virus remains trapped in the endosome and does not trigger an immune response.

Through biopanning using the entire virus as the antigen, antibodies specific to the E protein (the outer part of JEV) were successfully isolated. JEV envelope proteins are essential for viral attachment, fusion, virulence, cellular tropism, hemagglutination, and inducing protective immune responses [46]. Because of these vital functions, many antibodies have been developed to target JEV E proteins [44]. Like other flaviviral E proteins, the JEV E protein is divided into three domains. The most effective antibodies against flaviviruses typically bind to DIII, where the most unique epitopes are concentrated [14,46,47].

Based on in silico analysis, DE2 bound to three domains of the JEV E protein; therefore, this binding contributed to the scFv’s ability to neutralize JEV infection. D10 and C2 showed binding activity without neutralizing effects, hypothetically because their binding was confined to domains I and II. Several studies have shown that antibodies targeting domains I and II are less protective than those targeting domain III. [48–50]. Although DG2 is also a double-domain VL that binds to three domains, it only binds to a single amino acid in DIII. This supports the hypothesis that DG2 is unable to block JEV infection. Furthermore, DE2 is predicted to exhibit high specificity across all JEV genotypes but show no activity against other flavivirus families, such as dengue and Zika viruses. However, these binding site predictions still need to be confirmed by CRYO-EM or crystal structure analysis to determine the exact epitopes and paratopes [14,51].

In summary, through phage display and engineering, we obtained a double-domain light chain that can neutralize JEV. Neutralization occurred by blocking fusion with the endosomal membrane, preventing the JEV genome from being released into the cytoplasm. Based on an in silico study, this blocking was attributed to DE2 binding across all domains of the E protein, thereby interfering with the conformational change required for membrane fusion. Further animal protection studies are necessary to determine the neutralization capacity of this antibody.

## Acknowledgments

This work is supported by Korea Institute of Marine Science & Technology Promotion (KIMST) funded by the Ministry of Oceans and Fisheries, Korea (RS-2021-KS211475), and supported by the Education and Research Program for Future Human Resources in Applied Life Sciences, BK21 FOUR Program, funded by the Ministry of Education (MOE), Republic of Korea. The funders were not involved in study design, data collection and analysis, preparation, or the decision to submit the manuscript for publication.

## Data availability

The original contributions presented in the study are included in the article and Supplementary material; further inquiries can be directed to the corresponding authors.

## References

[1] Li C, Chen X, Hu J, Jiang D, Cai D, Li Y. A Recombinant Genotype I Japanese Encephalitis Virus Expressing a Gaussia Luciferase Gene for Antiviral Drug Screening Assay and Neutralizing Antibodies Detection. Int J Mol Sci 2022;23:15548. 10.3390/ijms232415548.

[2] Monath TP. Japanese Encephalitis: Risk of Emergence in the United States and the Resulting Impact. Viruses 2023;16:54. 10.3390/v16010054.

[3] Quan TM, Thao TTN, Duy NM, Nhat TM, Clapham H. Estimates of the global burden of Japanese encephalitis and the impact of vaccination from 2000-2015. Elife 2020;9. 10.7554/eLife.51027.

[4] Suresh KP, Nayak A, Dhanze H, Bhavya AP, Shivamallu C, Achar RR, et al. Prevalence of Japanese encephalitis (JE) virus in mosquitoes and animals of the Asian continent: A systematic review and meta-analysis. J Infect Public Health 2022;15:942–9. 10.1016/j.jiph.2022.07.010.

[5] Ladreyt H, Chevalier V, Durand B. Modelling Japanese encephalitis virus transmission dynamics and human exposure in a Cambodian rural multi-host system. PLoS Negl Trop Dis 2022;16:e0010572. 10.1371/journal.pntd.0010572.

[6] Sharma KB, Vrati S, Kalia M. Pathobiology of Japanese encephalitis virus infection. Mol Aspects Med 2021;81:100994. 10.1016/j.mam.2021.100994.

[7] Lee H-J, Choi H, Park KH, Jang Y, Hong Y, Kim YB. Retention of neutralizing antibodies to Japanese encephalitis vaccine in age groups above fifteen years in Korea. International Journal of Infectious Diseases 2020;100:53–8. 10.1016/j.ijid.2020.08.037.

[8] Shen K, Wang G, Yang H, Kang X, Yang L, Yuan Y, et al. Generation of soluble, immunoreactive recombinant JEV E protein through a simplified inclusion body extraction and refolding approach in vitro. Heliyon 2024;10:e34372. 10.1016/j.heliyon.2024.e34372.

[9] Gupta N, de Wispelaere M, Lecerf M, Kalia M, Scheel T, Vrati S, et al. Neutralization of Japanese Encephalitis Virus by heme-induced broadly reactive human monoclonal antibody. Sci Rep 2015;5:16248. 10.1038/srep16248.

[10] Kumar S, Verma A, Yadav P, Dubey SK, Azhar EI, Maitra SS, et al. Molecular pathogenesis of Japanese encephalitis and possible therapeutic strategies. Arch Virol 2022;167:1739–62. 10.1007/s00705-022-05481-z.

[11] Luca VC, AbiMansour J, Nelson CA, Fremont DH. Crystal Structure of the Japanese Encephalitis Virus Envelope Protein. J Virol 2012;86:2337–46. 10.1128/JVI.06072-11.

[12] Huang R, He Y, Zhang C, Luo Y, Chen C, Tan N, et al. The mutation of Japanese encephalitis virus envelope protein residue 389 attenuates viral neuroinvasiveness. Virol J 2024;21:128. 10.1186/s12985-024-02398-8.

[13] Wu K-P, Wu C-W, Tsao Y-P, Kuo T-W, Lou Y-C, Lin C-W, et al. Structural Basis of a Flavivirus Recognized by Its Neutralizing Antibody. Journal of Biological Chemistry 2003;278:46007–13. 10.1074/jbc.M307776200.

[14] Qiu X, Lei Y, Yang P, Gao Q, Wang N, Cao L, et al. Structural basis for neutralization of Japanese encephalitis virus by two potent therapeutic antibodies. Nat Microbiol 2018;3:287–94. 10.1038/s41564-017-0099-x.

[15] Nieri P, Donadio D, Rossi S, Adinolfi B, Podesta A. Antibodies for Therapeutic Uses and the Evolution of Biotechniques. Curr Med Chem 2009;16:753–79. 10.2174/092986709787458380.

[16] Emanuel P. Recombinant antibodies: a new reagent for biological agent detection. Biosens Bioelectron 2000;14:751–9. 10.1016/S0956-5663(99)00058-5.

[17] Monnier P, Vigouroux R, Tassew N. In Vivo Applications of Single Chain Fv (Variable Domain) (scFv) Fragments. Antibodies 2013;2:193–208. 10.3390/antib2020193.

[18] Ahmad ZA, Yeap SK, Ali AM, Ho WY, Alitheen NBM, Hamid M. scFv Antibody: Principles and Clinical Application. Clin Dev Immunol 2012;2012:1–15. 10.1155/2012/980250.

[19] Bazan J, Całkosiński I, Gamian A. Phage display—A powerful technique for immunotherapy. Hum Vaccin Immunother 2012;8:1817–28. 10.4161/hv.21703.

[20] Cho S-H, Kil E-J, Cho S, Byun H-S, Kang E-H, Choi H-S, et al. Development of novel detection system for sweet potato leaf curl virus using recombinant scFv. Sci Rep 2020;10:8039. 10.1038/s41598-020-64996-0.

[21] Safdari Y. Engineering of single chain antibodies for solubility. Int Immunopharmacol 2018;55:86–97. 10.1016/j.intimp.2017.11.046.

[22] Chen Y, Zhu X, Zhang X, Liu B, Huang L. Nanoparticles Modified With Tumor-targeting scFv Deliver siRNA and miRNA for Cancer Therapy. Molecular Therapy 2010;18:1650–6. 10.1038/mt.2010.136.

[23] Hoang PT, Luong QXT, Cho S, Lee Y, Na K, Ayun RQ, et al. Enhancing neutralizing activity against influenza H1N1/PR8 by engineering a single-domain VL-M2 specific into a bivalent form. PLoS One 2022;17:e0273934. 10.1371/journal.pone.0273934.

[24] Park Y, Kim A-R, Hwang Y-H, Yang H, Lee J-W, Kim MY, et al. Comparison of plaque reduction and focus reduction neutralization tests for the measurement of neutralizing antibody titers against japanese encephalitis virus. J Virol Methods 2022;306:114540. 10.1016/j.jviromet.2022.114540.

[25] Seo H, Lubis ADM, Lee S. A Novel Specific Single-Chain Variable Fragment Diagnostic System for Viral Hemorrhagic Septicemia Virus. Marine Biotechnology 2022;24:979–90. 10.1007/s10126-022-10161-9.

[26] de Wildt RMT, Mundy CR, Gorick BD, Tomlinson IM. Antibody arrays for high-throughput screening of antibody–antigen interactions. Nat Biotechnol 2000;18:989–94. 10.1038/79494.

[27] Hoang PT, Luong QXT, Ayun RQ, Lee Y, Oh K-J, Kim T, et al. A synergistic therapy against influenza virus A/H1N1/PR8 by a HA1 specific neutralizing single-domain VL and an RNA hydrolyzing scFv. Front Microbiol 2024;15. 10.3389/fmicb.2024.1355599.

[28] Luong QXT, Hoang PT, Lee Y, Ayun RQ, Na K, Park S, et al. An RNA-hydrolyzing recombinant minibody prevents both influenza A virus and coronavirus in co-infection models. Sci Rep 2024;14:8472. 10.1038/s41598-024-52810-0.

[29] Hong J-M, Munna AN, Moon J-H, Kim J-H, Seol J-W, Eo S-K, et al. Antiviral activity of prion protein against Japanese encephalitis virus infection in vitro and in vivo. Virus Res 2023;338:199249. 10.1016/j.virusres.2023.199249.

[30] Berry G, Tse L. Virus Binding and Internalization Assay for Adeno-associated Virus. Bio Protoc 2017;7. 10.21769/BioProtoc.2110.

[31] Gupta SK, Singh S, Nischal A, Pant KK, Seth PK. Molecular docking and simulation studies towards exploring antiviral compounds against envelope protein of Japanese encephalitis virus. Network Modeling Analysis in Health Informatics and Bioinformatics 2013;2:231–43. 10.1007/s13721-013-0040-z.

[32] Zhang H, Li D, Zheng J, Bao J, Wang Z, Qiu Y, et al. Sheep serve as amplifying hosts of Japanese encephalitis virus, increasing the risk of human infection. Sci Adv 2025;11. 10.1126/sciadv.ads7441.

[33] Srivastava KS, Jeswani V, Pal N, Bohra B, Vishwakarma V, Bapat AA, et al. Japanese Encephalitis Virus: An Update on the Potential Antivirals and Vaccines. Vaccines (Basel) 2023;11:742. 10.3390/vaccines11040742.

[34] Kitidee K, Samutpong A, Pakpian N, Wisitponchai T, Govitrapong P, Reiter RJ, et al. Antiviral effect of melatonin on Japanese encephalitis virus infection involves inhibition of neuronal apoptosis and neuroinflammation in SH-SY5Y cells. Sci Rep 2023;13:6063. 10.1038/s41598-023-33254-4.

[35] Chen I-C, Chiu Y-K, Yu C-M, Lee C-C, Tung C-P, Tsou Y-L, et al. High throughput discovery of influenza virus neutralizing antibodies from phage-displayed synthetic antibody libraries. Sci Rep 2017;7:14455. 10.1038/s41598-017-14823-w.

[36] van Dorsten RT, Lambson BE, Wibmer CK, Weinberg MS, Moore PL, Morris L. Neutralization Breadth and Potency of Single-Chain Variable Fragments Derived from Broadly Neutralizing Antibodies Targeting Multiple Epitopes on the HIV-1 Envelope. J Virol 2020;94. 10.1128/JVI.01533-19.

[37] Yu K, Liu B, Yu H, Sun C, Wang X, Li G, et al. A neutralizing bispecific single-chain antibody against SARS-CoV-2 Omicron variant produced based on CR3022. Front Cell Infect Microbiol 2023;13. 10.3389/fcimb.2023.1155293.

[38] Kim DY, Hussack G, Kandalaft H, Tanha J. Mutational approaches to improve the biophysical properties of human single-domain antibodies. Biochimica et Biophysica Acta (BBA) - Proteins and Proteomics 2014;1844:1983–2001. 10.1016/j.bbapap.2014.07.008.

[39] Henry KA, Kim DY, Kandalaft H, Lowden MJ, Yang Q, Schrag JD, et al. Stability-Diversity Tradeoffs Impose Fundamental Constraints on Selection of Synthetic Human VH/VL Single-Domain Antibodies from In Vitro Display Libraries. Front Immunol 2017;8. 10.3389/fimmu.2017.01759.

[40] Hoang PT, Luong QXT, Cho S, Lee Y, Na K, Ayun RQ, et al. Enhancing neutralizing activity against influenza H1N1/PR8 by engineering a single-domain VL-M2 specific into a bivalent form. PLoS One 2022;17:e0273934. 10.1371/journal.pone.0273934.

[41] Miller BR, Demarest SJ, Lugovskoy A, Huang F, Wu X, Snyder WB, et al. Stability engineering of scFvs for the development of bispecific and multivalent antibodies. Protein Engineering, Design and Selection 2010;23:549–57. 10.1093/protein/gzq028.

[42] Kang TH, Seong BL. Solubility, Stability, and Avidity of Recombinant Antibody Fragments Expressed in Microorganisms. Front Microbiol 2020;11. 10.3389/fmicb.2020.01927.

[43] Wang Y, Yuan W, Guo S, Li Q, Chen X, Li C, et al. A 33-residue peptide tag increases solubility and stability of Escherichia coli produced single-chain antibody fragments. Nat Commun 2022;13:4614. 10.1038/s41467-022-32423-9.

[44] Zhu Y, He Z, Qi Z. Virus-host Interactions in Early Japanese Encephalitis Virus Infection. Virus Res 2023;331:199120. 10.1016/j.virusres.2023.199120.

[45] Safdari Y, Ahmadzadeh V, Khalili M, Jaliani HZ, Zarei V, Erfani-Moghadam V. Use of Single-Chain Antibody Derivatives for Targeted Drug Delivery. Molecular Medicine 2016;22:258–70. 10.2119/molmed.2016.00043.

[46] Lin C-W, Wu S-C. A Functional Epitope Determinant on Domain III of the Japanese Encephalitis Virus Envelope Protein Interacted with Neutralizing-Antibody Combining Sites. J Virol 2003;77:2600–6. 10.1128/JVI.77.4.2600-2606.2003.

[47] Islam MdD, Islam MM, Inoue A, Yesmin S, Brindha S, Yoshizue T, et al. Neutralizing antibodies against the Japanese encephalitis virus are produced by a 12 kDa E. coli-expressed envelope protein domain III (EDIII) tagged with a solubility-controlling peptide. Vaccine 2025;56:127143. 10.1016/j.vaccine.2025.127143.

[48] Smith SA, de Alwis AR, Kose N, Harris E, Ibarra KD, Kahle KM, et al. The Potent and Broadly Neutralizing Human Dengue Virus-Specific Monoclonal Antibody 1C19 Reveals a Unique Cross-Reactive Epitope on the bc Loop of Domain II of the Envelope Protein. MBio 2013;4. 10.1128/mBio.00873-13.

[49] Oliphant T, Nybakken GE, Engle M, Xu Q, Nelson CA, Sukupolvi-Petty S, et al. Antibody Recognition and Neutralization Determinants on Domains I and II of West Nile Virus Envelope Protein. J Virol 2006;80:12149–59. 10.1128/JVI.01732-06.

[50] Pierson TC, Diamond MS. Molecular mechanisms of antibody-mediated neutralisation of flavivirus infection. Expert Rev Mol Med 2008;10:e12. 10.1017/S1462399408000665.

[51] Nybakken GE, Oliphant T, Johnson S, Burke S, Diamond MS, Fremont DH. Structural basis of West Nile virus neutralization by a therapeutic antibody. Nature 2005;437:764–9. 10.1038/nature03956

